# Oral pyruvate prevents glaucomatous neurodegeneration

**DOI:** 10.1101/2020.05.02.072215

**Authors:** Pete A Williams, Jeffrey M Harder, Chelsea Guymer, John P M Wood, Evangelia Daskalaki, Glyn Chidlow, Brynn H Cardozo, Nicole E Foxworth, Kelly E Cochran, Tionna B Ouellette, Craig E Wheelock, Robert J Casson, Simon W M John

## Abstract

Intraocular pressure-sensitive retinal ganglion cell degeneration is a hallmark of glaucoma, the leading cause of irreversible blindness. Converging evidence indicates that age-related bioenergetic insufficiency increases the vulnerability of retinal ganglion cells to intraocular pressure. To investigate further, we used metabolomics and RNA-sequencing to examine early glaucoma in DBA/2J mice. We demonstrate an intraocular pressure-dependent decline in retinal pyruvate levels coupled to dysregulated glucose metabolism prior to detectable optic nerve degeneration. Oral supplementation of pyruvate strongly protected from neurodegeneration in pre-clinical models of glaucoma. We detected mTOR activation at the mechanistic nexus of neurodegeneration and metabolism. Rapamycin-induced inhibition of mTOR robustly prevented glaucomatous neurodegeneration. Bioenergetic enhancement, in combination with intraocular pressure reduction, therefore provides a readily translatable strategy that warrants investigation in clinical trials.

**Funding:** Vetenskapsrådet 2018-02124 and StratNeuro StartUp grant (PAW). Pete Williams is supported by the Karolinska Institutet in the form of a Board of Research Faculty Funded Career Position and by St. Erik Eye Hospital philanthropic donations. EY011721 and the Barbra and Joseph Cohen Foundation and startup funds from Columbia University (SWMJ). Simon John is an Investigator of HHMI.

## Main Text

Glaucoma is characterized by the progressive dysfunction and subsequent degeneration of retinal ganglion cells (RGCs) and their axons in the optic nerve. Increasing evidence suggests that risk factors for glaucoma conspire to reduce energy production in the retina and optic nerve. Age- and intraocular pressure-dependent metabolic deficits are emerging as a critical factor influencing RGC vulnerability in glaucoma (*1, 2*). Here, we demonstrate that elevated intraocular pressure (IOP) alters metabolism beyond the changes caused by aging by decreasing retinal pyruvate levels. Pyruvate is an important nutrient for diverging energy metabolism pathways and for balancing energy needs in different physiological contexts (*3, 4*).

To further understand how elevated IOP alters energy metabolism in RGCs in a chronic pre-clinical model of glaucoma, we analyzed RNA-sequencing datasets from RGCs of 9 month-old DBA/2J (D2) and D2-*Gpnmb*^+^ mice (a strain-matched non-glaucoma control). At 9 months of age in our colony, D2 eyes have had ongoing high IOP, but have not yet developed glaucoma, allowing for the assessment of early pre-degenerative molecular changes. We focused on changes in energy metabolism defined by functional analysis of differentially expressed genes. This identified many early transcriptional changes to *glycolysis* (produces pyruvate), *mannose and fructose metabolism, gluconeogenesis* (utilizes pyruvate), and *pyruvate metabolism* in general. Higher expression of pertinent regulatory genes also occurred in D2 RGCs, including the glucose transporter *Glut3*, the glycolytic regulatory gene *Pfkfb3*, pyruvate transporters *Mpc1* and *Mpc2*, and pyruvate dehydrogenase component *Pdha1* (**Fig S1A**). Given the theme of altered pyruvate metabolism in the RGC gene expression data, we next determined how retinal pyruvate levels were affected. Pyruvate levels declined with increasing exposure to elevated IOP in D2 retinas, but did not change significantly with age alone (in control D2-*Gpnmb*^+^ mice) (**Fig S1B**). Thus, elevated IOP has a significant impact on pyruvate production or metabolism. These data are consistent with a genetic program in RGCs favoring pyruvate use in response to chronically elevated IOP; or could indicate a loss in pyruvate production during IOP-related stress (or a combination of both). Since pyruvate is a key nutrient in various energy metabolism pathways, pyruvate depletion may increase RGC dysfunction and vulnerability. Immuno-labeling for mitochondrial pyruvate carriers, MPC1 and MPC2, confirmed that RGCs have the machinery to direct pyruvate into the citric acid cycle. These proteins were present in high levels in RGCs across a range of species (mouse, rat, primate, human) highlighting the potential importance of pyruvate metabolism to RGCs (**Fig S1C and D**).

In addition to a major role in energy metabolism, pyruvate has been shown to protect against PARP activation, oxidative stress, and neuroinflammation (*4, 5*), all of which have been associated with glaucoma (*6-9*). We tested the ability of pyruvate to protect stressed RGCs utilizing cultured retinal neurons and assessing the effects of pyruvate treatment following glucose deprivation or oxidative stress. In both models, pyruvate supplementation enhanced neuronal survival (**Fig S2**). Inhibiting monocarboxylate transporters, which facilitate pyruvate uptake into cells, with 4CIN potently inhibited the protective action of pyruvate (**Fig S2**). This suggests that pyruvate is an important energy source and acts as a protectant for RGCs under stress. The neuroprotective effects of pyruvate were also assessed in a model of axon injury that leads to programmed cell death similar to glaucoma. In mouse retina explant cultures pyruvate supplementation prevented degeneration of axotomized RGCs (**Fig S3**).

We further assessed the efficacy of long-term oral pyruvate supplementation *in vivo*. Pyruvate-treated D2 mice showed increased levels of retinal pyruvate at 9 months, an age prior to histologically detectable glaucoma (**Fig 1A, Fig S4**). Continued treatment reduced the onset of optic nerve degeneration and prevented RGC loss at older ages, yielding a ∼2-fold decrease in the incidence of glaucoma (**Fig 1B-E**). RGCs of aged, pyruvate-treated D2 mice had enhanced visual function compared to untreated mice (as assessed by pattern electroretinography; PERG, a sensitive measure of RGC function in humans and animals) and had intact, long-range axon projections capable of anterograde axoplasmic transport (**Fig 1F-I**). As acute or sub-acute high IOP insults contribute to clinical incidences of glaucoma, we also tested pyruvate in an, inducible model of glaucoma. In this model, laser photocoagulation of limbal blood vessels in rats induced a sub-acute elevation of IOP which leads to a significant loss of RGCs. Pyruvate treatment increased RGC survival, decreased RGC axon cytoskeletal damage and loss, and decreased microglial activation (**Fig 1J-O**). Thus, pyruvate treatment has promising potential to reduce glaucoma in patients with diverse IOP insults.

**Fig. 1.**
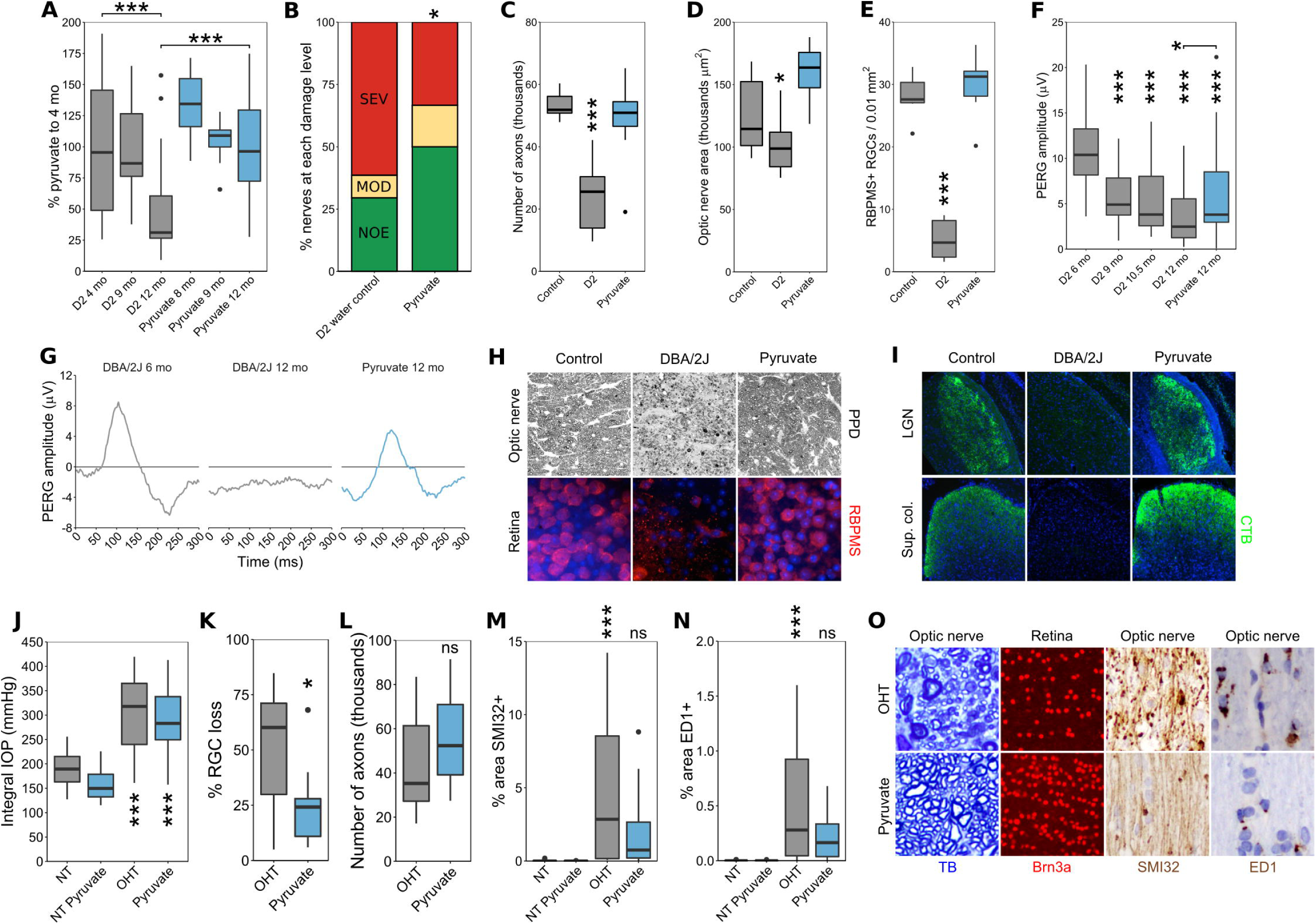
Pyruvate protects from glaucomatous neurodegeneration in chronic mouse and rat models. (**A-I**) DBA/2J mice. (**A**) Retinal pyruvate levels decline following the onset of elevated IOP and are recovered by oral pyruvate treatment (*n* > 24/group). (**B and H**) Pyruvate intervention protected from optic nerve degeneration as assessed by PPD staining (paraphenylenediamine; a sensitive stain for damaged axons) in D2 mice (*n* = 60/group). Green, no or early damage [<5% axon loss; no or early (NOE)]; yellow, moderate damage (∼30% axon loss; MOD); red, severe (>50% axon loss; SEV) damage. (**C**) Counts and (**D**) optic nerve areas from D2 mice demonstrate a robust axon protection by pyruvate treatment (*n* = 10/group). (**E and H**) Pyruvate protected from RGC soma loss (RBPMS is a specific marker of RGCs in the retina; *n* = 6/group). (**F and G**) Pyruvate rescued PERG amplitude so that 12 mo old treated mice were not significantly different to 9 mo untreated mice that had not yet developed glaucomatous degeneration (PERG is a sensitive measure of RGC function in animals and humans; *n* > 20/group). Individual example tracings are shown in **G**. (**I**) Pyruvate protects from the loss of anterograde axoplasmic transport as assessed by CT-β tracing to the dLGN (dorsal lateral geniculate nucleus) and SC (superior colliculus) (*n* = 6/group). (**J-O**) Inducible rat model. (**J**) Laser photocoagulation of limbal blood vessels in rats caused a sub-acute elevation of IOP (ocular hypertension; OHT). The magnitude of IOP elevation is unchanged by pyruvate treatment. (**K and L**) Pyruvate treatment in the rat prevents RGC soma loss and axon loss in the optic nerve. Examples are shown in **O**. Brn3a is a specific marker of RGCs in the rat retina. (**M-O**) Pyruvate treatment decreases RGC axon cytoskeletal damage (SMI32+ dystrophic axons) and decreases microglia activation (ED1+ microglia) (*n* = 18/group for **J-O**).

To better understand the metabolic changes in glaucoma, we performed metabolomics on single retinas at time points matching our RNA-sequencing analysis. We analyzed D2-*Gpnmb*^+^ control mice, untreated D2 mice, and D2 mice treated with pyruvate or nicotinamide (another metabolite we have demonstrated to be strongly neuroprotective (*10*)). Eighty-six metabolites were identified at Level 1 accuracy according to the Metabolomics Standards Initiative (except for isoleucine) (*11*), the majority of which were altered following elevated IOP (39 increased, 9 decreased in glaucoma compared to controls) (**Fig 2A-C, Fig S5, Table S1**). Pathway analyses of the altered metabolites showed enrichment in pathways related to glucose metabolism and oxidative stress, indicating a preponderance in changes related to energy metabolism. Similar pathways were enriched in gene expression changes specific to RGCs (**Fig 2D-F, Table S2**). These data suggest during ocular hypertensive stress there is metabolic-coupling between RGCs and other retinal cell types and/or that the observed changes in metabolites occur in RGCs. The largest change in a single metabolite following elevated IOP was glucose (52-fold increase). The increased glucose may reflect altered glycolysis or other features of glucose metabolism including conversion of astrocytic glycogen to support an increased energy demand due to glaucoma (*2*). We previously demonstrated PARP upregulation in RGCs during D2 glaucoma and PARP can inhibit glycolysis by impacting hexokinase (*5*). Elevated glucose is associated with poor outcomes in brain injury and disease (*12, 13*), and high glucose concentrations lead to decreased mitochondrial cristae and mitochondrial remodeling in diabetic and cell models (*14, 15*). D2 RGC mitochondria have decreased cristae at corresponding disease time points (*10*). D2 mice do not have hyperglycemia (*16, 17*), implying that retinal glucose metabolism itself is disrupted.

**Fig. 2.**
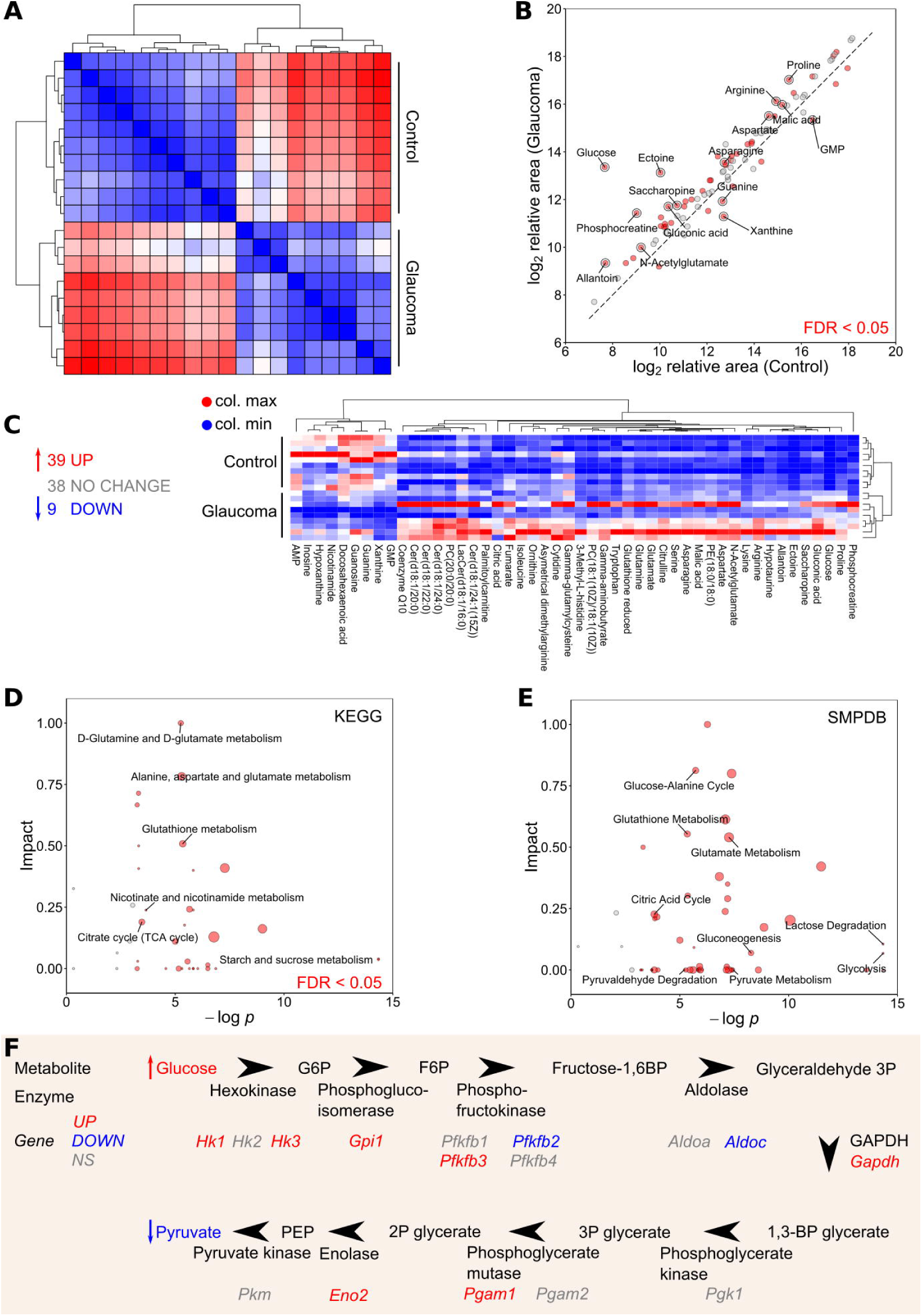
Retinal metabolic dysfunction occurs early following the onset of elevated IOP in D2 mice. (**A**) Whole retina samples underwent metabolomic profiling (at 9 mo of age, *n* = 10/group, control = D2-*Gpnmb*^+^, glaucoma = D2). HC (*Spearman’s*) defined metabolically distinct groups. (**B and C**) 86 metabolites were conclusively identified with Level 1 accuracy according to the Metabolomics Standards Initiative (*11*) (except for isoleucine) (39 increased, 9 decreased in D2 pre-glaucomatous retinas compared to controls). All metabolites are shown in **B**, significantly changed metabolites are shown in **C**. (**D**) KEGG and (**E**) SMPDB pathway analysis of metabolites demonstrates significant pathway enrichment of pathways related to glucose and pyruvate metabolism and NAD metabolism. (**F**) A schematic pathway of glycolysis integrating metabolomic and RNA-sequencing data.

An additional key difference in metabolic profiles between control and glaucoma retinas was a decrease in retinal nicotinamide (NAM). NAM levels were lower in D2 retinas compared to controls consistent with decreased retinal NAD levels in this model (*10*) and decreased serum NAM levels in primary open angle glaucoma patients (*18*). We have previously demonstrated that NAM confers a robust neuroprotection against damaging IOP insults. NAM supplementation prevents age-dependent decreases in NAD levels, transcriptional reprogramming of RGCs, mitochondrial decline, and RGC neurodegeneration (*1, 10, 19, 20*). Unexpectedly, our metabolomic analysis of retinas from NAM treated and untreated D2 mice did not detect differences outside of NAM itself (*data not shown*). Pyruvate treatment rescued more IOP-dependent changes in the retinal metabolome than NAM. Pyruvate treated D2 mice had differences in 27 metabolites compared to untreated mice, including lower levels of glucose and higher levels of NAM (**Fig S6**). The increased NAM most likely reflects increased NAD availability to NAD-consuming enzymes that generate NAM in pyruvate-treated mice. Thus, in addition to providing an energy substrate for mitochondrial respiration, pyruvate treatment affected NAD use. As NAD is a critical factor for glycolysis, together a decrease in NAD and pyruvate, and increase in glucose, are consistent with a glycolytic deficiency. During D2 glaucoma, RGC axons were recently shown to have reduced capacity to increase glycolysis in response to metabolic demand (*21*). Although the regulation of glycolysis is complex, glycolytic insufficiency can occur if NAD levels decrease as the reduction of NAD^+^ to NADH is essential for conversion of glyceraldehyde-3-phosphate to 1,3-bisphosphoglycerate. Additionally, pyruvate metabolism is important for replacing NAD^+^ (*3*). Thus, the reduced NAD and pyruvate levels are likely to contribute to reduced glycolytic capacity in D2 RGCs, and *vice versa*.

Pyruvate and NAM are ideal treatments for clinical use, with long histories and favorable safety profiles in humans (*1, 22*). The most protective dose of NAM that we previously demonstrated in D2 mice is large, complicating compliance, and may be unachievable or unsafe in some patients (*1, 10*). Importantly, our metabolomic data suggested that pyruvate and NAM treatments may be complementary. Thus we hypothesized that supplementing the diet of D2 mice with both pyruvate and a suboptimal dose of NAM may be beneficial. Combination therapy of pyruvate and NAM lowered the risk of RGC loss and optic nerve degeneration in D2 mice by ∼2.6 fold, more than either treatment regime alone (**Fig S7**). This combination therapy showed that targeting IOP-dependent (declining pyruvate) and age-related (declining NAD) changes in glaucoma can be additive and might represent an ideal combination for human translation.

These metabolic and treatment data suggest that dysfunctional metabolism may underlie RGC vulnerability in glaucoma. Related to this, pathway analysis of our RNA-sequencing data predicted activation of the mTOR signaling pathway in RGCs in ocular hypertensive eyes (**Fig 3A and B**). mTOR is a master regulator of metabolism, and mTOR complex 1 (mTORC1) modulates glucose use and glycolysis through *Hif1*α (*23*). Hif1α is increased in the retinas of D2 mice following high IOP (*10*) and D2 optic nerves have a decreased maximal respiratory rate and altered glycolytic responses (*21*). Analysis of retinal proteins by an mTOR pathway antibody array demonstrated a specific increase in phospho-S6 and phospho-S6K, classic indicators of mTOR activation (**Fig 3C**). The observed increase in mTOR pathway activity corresponded with PARP activation (*10*), and increased levels of glucose and related metabolites. Together, these data collectively support an increase in neuronal energy use following IOP-related stress. As mTOR inhibition can protect neurons from cell death by reducing energy use (*24*), we tested the role of mTOR in D2 mice. D2 mice were treated with rapamycin to inhibit mTOR-dependent metabolic reprogramming. Rapamycin is an mTOR inhibitor that promotes survival mechanisms associated with fasting and decreased energy use (*23*). Providing evidence that mTOR and imbalanced energy regulation represent a critical target for therapies, rapamycin treatment alone led to a ∼2.6-fold decrease in the incidence of severe glaucoma in D2 mice (**Fig 3D-F, Fig S4**).

**Fig. 3.**
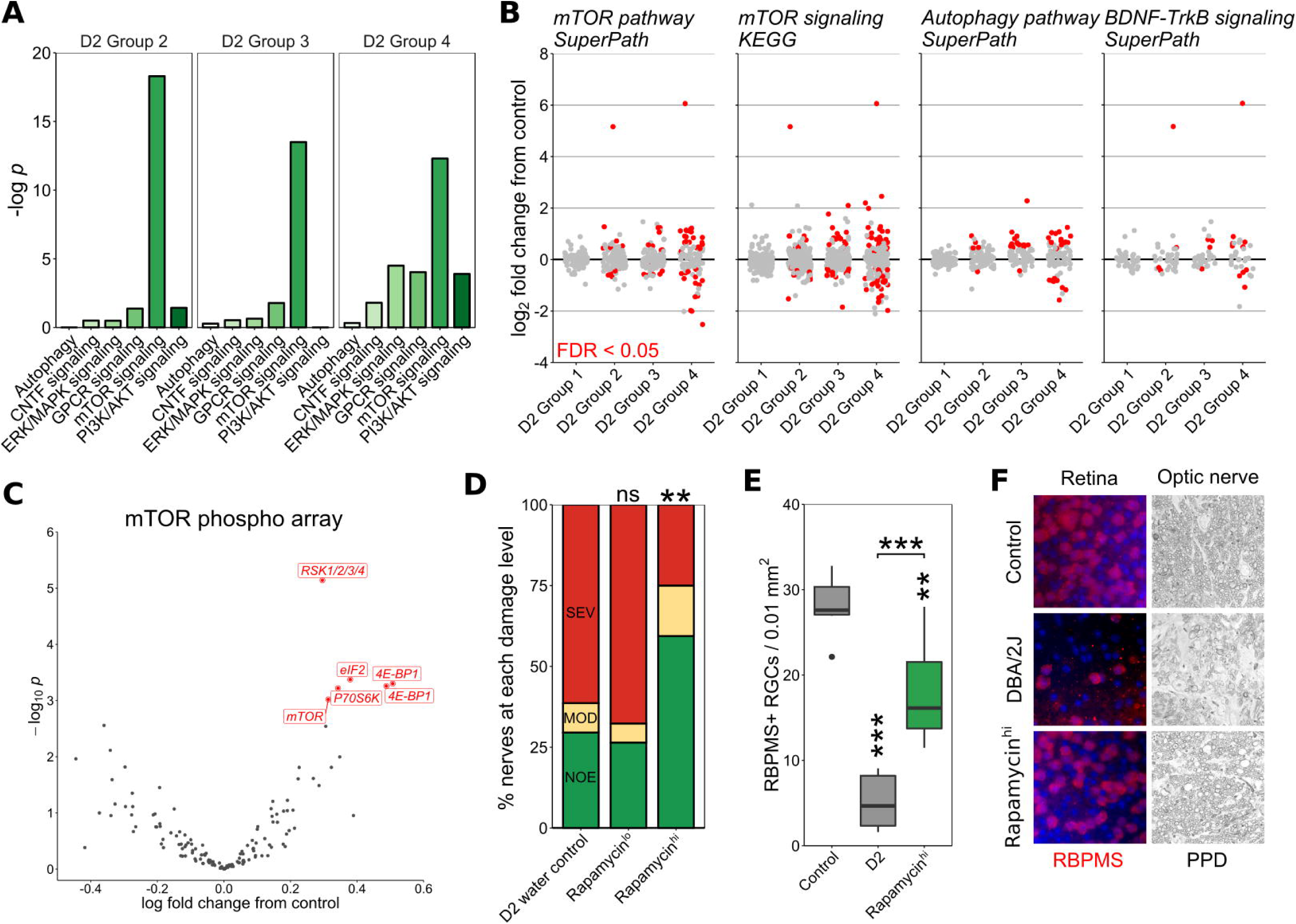
mTOR activation occurs early in glaucoma with rapamycin treatment protecting from glaucoma. (**A-B**) Pathway analysis of RNA-sequencing data from pre-glaucomatous D2 RGCs demonstrates significant enrichment of mTOR-related metabolic pathways. D2 Groups 1-4 represent increasing disease-status at the transcriptomic level as identified in Williams *et al*. 2017 (*10*). Group 1 is indistinguishable from no glaucoma controls with no significant pathway differences. (**C**) mTOR pathway antibody array demonstrated a specific increase in phospho-S6 and phospho-S6K, classic markers of mTOR activation. The two labels for 4E-BP1 are separate phospho-sites, Thr 36 and Thr 45. (**D-F**) Rapamycin intervention protected from optic nerve degeneration (PPD staining) and RGC soma loss (RBPMS labeling).

In conclusion, these findings are consistent with a model of glaucoma where elevated IOP disrupts energy homeostasis by affecting the availability of metabolic energy substrates. Further affected by low availability of coenzymes, such as NAD^+^, RGCs ultimately lack the energy needed to function and cope with stress and inflammation associated with age and ocular hypertension. Energy supplements not only reduce degeneration, but also improve visual function. In this light, decreased visual function prior to degeneration may be an adaptive response to reduce energy consumption and signal a need for energy supplements. Although the combination of decreased NAD and pyruvate represent a critical component of RGC susceptibility in glaucoma, it is directly therapeutically approachable. Combining vitamin and energy supplements with established IOP-lowering therapies represents a powerful therapeutic strategy for human glaucoma that necessitates clinical investigation.

## Acknowledgements

The Authors would like to thank Mimi de Vries and Amy Bell for assistance with organizing mouse colonies and intraocular pressure measurements, and the staff of the histology, gene expression services, and computational sciences at The Jackson Laboratory. All data presented is available in the supplementary materials.

## Materials and methods

### Mouse strains, breeding and husbandry

Mice were housed in a 14h light / 10h dark cycle with food and water available ad libitum as previously reported (*10*). All breeding and experimental procedures were undertaken in accordance with the Association for research for Vision and Ophthalmology Statement for the Use of Animals in Ophthalmic Research. The Institutional Biosafety Committee and the Animal Care and Use Committee at The Jackson Laboratory approved this study. C57BL/6J (B6), DBA/2J (D2; glaucoma), and DBA/2J-*Gpnmb*^*R150X*^ (D2-*Gpnmb*^+^; control) strains were utilized. For aged glaucoma experiments mice were administered nicotinamide (NAM) or pyruvate in water starting at 6 mo (prophylactic, prior to IOP elevation in almost all eyes) or 9 mo (when the majority of eyes have high IOP and molecular changes, but no detectable neurodegeneration). NAM (550 mg/kg/d, PanReac AppliChem) and/or pyruvate (500 mg/kg/d, Sigma) was dissolved in regular acid drinking water (350 ml) and changed once per week. Whole pens (*n* of mice = 5) were randomized between treatments and control. Rapamycin (Sigma) was administered in drinking water at doses of 0.5 mg/kg/d and 2 mg/kg/d.

### Clinical phenotyping

IOP elevation in D2 mice is subsequent to a pigment dispersing iris disease. In all experiments, the progression of the iris disease and intraocular pressure in mutant or drug-treated mice was compared to control D2 mice as previously described (*25*). In each experiment, iris disease and intraocular pressure were assessed. Iris disease was assessed at two-month intervals starting at 6 months of age until experiment completion. Intraocular pressure was measured at 45-day intervals beginning at 8.5-9 mo until experiment completion.

### Rat husbandry and experimental rat glaucoma models

Rat projects were approved by the Animal Ethics Committee of the SA Pathology / Central Adelaide Local Health Network (CALHN), Adelaide, and the University of Adelaide, and conforms to the 2004 Australian Code of Practice for the Care and Use of Animals for Scientific Purposes. Adult Sprague-Dawley rats (aged >10 weeks, >230g) were housed in a temperature and humidity controlled room with 12-hour light and dark cycles. Food and water were provided *ad libitum*. Rats were randomly assigned into control (water only) and pyruvate supplementation (500 mg/kg/day in normal drinking water) groups. Rats were commenced on pyruvate supplementation exactly one week prior to glaucoma induction and continued on pyruvate throughout the experiment. Animals were excluded from further analysis if severe hyphema developed subsequent to laser treatment (29% collectively, which was lower than the hyphema rate of 40% reported Levkovitch-Verbin *et al*. (*26*).

Experimental glaucoma was induced at day 0 in the right eye of all animals, leaving the left eye untouched to serve as a control. A slightly modified protocol of the method described by Levkovitch-Verbin *et al*. was used to induce ocular hypertension in the right eye of each animal by laser photocoagulation of the episcleral and circumlimbal vessels (*26*). In brief, 135-150 spots of 100 μm diameter, 340 mW power and 0.6 second duration were applied to the circumlimbal vessels. An additional 35-50 spots, 200 μm diameter, 300 mW power for 0.6 seconds duration, were delivered to the dominant superior, inferior and temporal episcleral veins whilst leaving the nasal episcleral vein and intersecting nasal margins of the circumlimbal vessel patent. Intraocular pressure was monitored using a rebound tonometer (TonoLab, Icare, Finland).

### Pattern electroretinography (PERG)

PERG was recorded subcutaneously from the snout of mice as previously reported (*10, 27*). Waveforms were retrieved using an asynchronous averaging method (*28*).

### Optic nerve assessment and determination of glaucomatous damage in DBA/2J mice

The processing of optic nerves and staining with paraphenylenediamine (PPD) was as published (*29*). A segment of optic nerve from within a region up to 1 mm from the posterior surface of the sclera was sectioned (1 µm thick sections) and stained with PPD. Typically 30-50 sections are taken from each nerve. Multiple sections of each nerve were considered when determining damage level. Optic nerves were analyzed and determined to have one of 3 damage levels; NOE (no or early damage), MOD (moderate damage), or SEV (severe damage) (*10*). Optic nerve axon counts were performed using AxonJ (*30*).

### Optic nerve assessment in a rat model of experimental glaucoma

A short piece of proximal optic nerve from the right treated eye of each rat, 1.5 mm behind the globe, was removed for resin processing. In short, proximal optic nerves were fixed by immersion in 2.5% glutaraldehyde with 4% paraformaldehyde in 0.1 M phosphate buffer, pH 7.4 for at least 24 hours. Optic nerves were then placed in osmium tetroxide in saline overnight and washed with cocadylate buffer at room temperature. Optic nerves were subsequently dehydrated in graded alcohols and embedded in epoxy resin for transverse sectioning. An ultramicrotome was used to cut sections at 1 μm, which were then mounted on glass slides, and enhanced with osmium tetroxide-induced myelin staining using 1% toluidine blue.

### Anterograde axon transport

Mice were anaesthetized using ketamine / xylazine and intraviteally injected with 2 μl AF488 cholera toxin subunit B (1 mg/ml in PBS) (ThermoFisher Scientific). After 72 h mice were anaesthetized and euthanized via 4% PFA cardiac perfusion. Brains and eyes were post-fixed in 4% PFA for an additional 24 h, cryoprotected in 30% sucrose in PBS overnight, OCT cryoembedded, and sectioned at 20 μm. AF488 was visualized using a Zeiss AxioObserver.

### Histology: whole-mounted retina

For immunofluorescence staining, mice were euthanized by cervical dislocation, their eyes enucleated and placed in 4% PFA ON. Retinas were dissected and whole-mounted onto slides, permeabilized with 0.1% Triton-X for 15 mins, blocked with 2% BSA in PBS and incubated overnight at RT in primary antibody (see **Table S2** for a complete list of antibodies used in this manuscript). After primary antibody incubation, retinas were washed 5 times in PBS, stained for 4 h at RT with secondary antibody. Slides were then washed a further 5 times with PBS, labelled with 500 ng/ml DAPI for 15 mins, mounted with fluoromount and sealed with nail-polish. For retinal sections, eyes were cryoprotected in 30% sucrose overnight, frozen in OCT, and cryosectioned at 18 μm. Slides were warmed to room temperature and the procedure above was followed. Retinas were imaged on a Zeiss AxioObserver for low resolution counts.

### Histology: retinal sections

Tissue sections from mouse, rat, marmoset, and human donors were used to visualize MPC expression. These studies were approved by the SA Pathology/CALHN Animal Ethics Committee (Adelaide, Australia), and conformed with the Australian Code of Practice for the Care and Use of Animals for Scientific Purposes, 2013, and with the ARVO Statement for the use of animals in vision and ophthalmic research. Ocular tissue was collected from adult marmosets (*Callithrix jacchus*), aged 10 to 14 years, belonging to the colony housed at the Queen Elizabeth Hospital (South Australia, Australia) that were being euthanized. Human ocular tissue for analysis was obtained from the Eye-Bank of South Australia, Flinders Medical Centre (Adelaide, Australia) following the guidelines of the Southern Adelaide Clinical Human Research Ethics Committee; all had been screened to ensure there was no underlying ocular disease and all were from Caucasian donors aged 50-65. There was no consistent orientation of globes with regard to the nasal-temporal, superior-inferior quadrants of the retina analyzed. Sections were typically taken at the level of the optic nerve head and hence comprise at least two quadrants. Globes that were used for immunohistochemistry were immersion-fixed in 10% buffered formalin or Davidson’s solution for 24 h (rat, mouse) or 48 h (for marmoset, human eyes), and transferred to 70% ethanol until processing. Eyes were then processed for routine paraffin-embedded sections. Globes were embedded sagittally. In all cases, 4 μm sections were cut. Colorimetric immunohistochemistry was performed as previously described (*31*). In brief, tissue sections were deparaffinised, endogenous peroxidase activity was blocked and high-temperature antigen retrieval was performed. Subsequently, sections were incubated in primary antibody, followed by consecutive incubations with biotinylated secondary antibody and streptavidin-peroxidase conjugate. Color development was achieved using 3,3’-diaminobenzidine. Confirmation of the specificity of antibody labeling was judged by the morphology and distribution of the labeled cells, by the absence of signal when the primary antibody was replaced by isotype/serum controls, and by the presence of a comparable signal when an alternative primary antibody was used.

### Mixed retinal cell culture

Sprague-Dawley rats were housed according to the Australian Code of Practice for the Care and Use of Animals for Scientific Purposes 2004, and the ARVO Statement for the Use of Animals in Ophthalmic and Vision Research. From stocks, litters of pups (1-3 days post-partum) were obtained to derive mixed retinal cell cultures. Cultures were prepared via a sequential trypsin- and mechanical-digest procedure and comprised neurons, including RGCs, glia, and photoreceptors, as previously described (*32*). Isolated cells were dispensed at 0.5 × 106 cells/ml onto 13 mm borosilicate glass coverslips (pre-coated for 15 minutes with 10 µg/ml poly-L-lysine), for immunocytochemistry, or into 96-well plates (CellPlus positive-charge-coated plates, Sarstedt), for viability assays. Cultures were routinely maintained under saturating humidity at 37°C in Minimal Essential Medium containing 10% (v/v) foetal bovine serum (FBS), 5 mM D-glucose, 2 mM L-glutamine, and penicillin/streptomycin.

For treatment of cell cultures, treatments were commenced at 7 days *in vitro* and were carried out for 24 h. For nutrient deprivation (ND) experiments, culture medium was replaced with one lacking FBS, glucose, pyruvate, and glutamine. Pyruvate was added at appropriate test concentrations (100 µM, 1 mM, 5 mM) and the monocarboxylate transport inhibitor, α-cyano-4-hydroxycinnamic acid (4CIN), was applied at 10 µM. For application of oxidative stress to cells, medium lacking FBS, glucose and pyruvate but containing 2 mM L-glutamine was used. Glutamine was present in this case because removal of all nutrients caused catastrophic death of all cells within 2-4 hours and this amino acid has been shown to support neuron survival *in vitro* (*33*). When cultures were to be used for immunocytochemistry, cells on coverslips were fixed for 10 minutes with 10% (w/v) neutral buffered formalin containing 1% (v/v) methanol. For viability assessment using the 3-(4,5-dimethylthiazol-2-yl)-2,5-diphenyltetrazolium bromide (MTT) assay, for the last hour of the incubation, medium and test compounds were removed and new medium free of potentially-confounding dead cells/cell debris was applied along with 0.5 mg/ml MTT. After one hour, medium was removed from wells again and remaining cells solubilized with dimethyl sulphoxide (DMSO) before colorimetric absorbance was determined at 570 nm (with 630 nm reference).

### Immunocytochemical assessment of mixed retinal cultures

Fixed cells were permeabilised in 0.1% Triton X-100 (v/v) in PBS for 15 minutes and then blocked in 3.3% (v/v) horse serum in PBS (PBS-HS). Primary antibodies were appropriately diluted in PBS-HS and applied to coverslips overnight at room temperature in a moist chamber. After overnight incubation, coverslips were washed in PBS and then labelling was completed by successive incubations with appropriate biotinylated secondary antibody (1:250) and fluorescent AlexaFluor-conjugated (488 or 594) streptavidin (1:500). Nuclear counterstaining was achieved with a 5 minute incubation between PBS washes with 500 ng/ml DAPI. In the case of double-labelling of cultures by two antibodies, one was developed as mentioned and the other, concurrently, with an appropriate secondary species-specific antibody directly linked to the opposite colored Alexafluor fluorescent label. Quantification of immunocytochemistry was achieved by manually counting labelled cells for each antibody in five fields per coverslip, averaging and rounding to the nearest integer; this represented a single determination.

### Whole retina explant culture

Whole retina explant culture was performed as previously described (*10*). Retinas were incubated in pyruvate in retinal explant media at the doses indicated. Retinas were incubated in 6-well plates at 37 °C and 4% CO_2_ for 5 days before being fixed in 4% PFA and stained with DAPI. Retinas were imaged on a Zeiss AxioObserver.

### Retinal pyruvate level quantification

For pyruvate quantification mice were euthanized by cervical dislocation and retinas were homogenized in ice-cold HBSS and pyruvate measured following the manufacturer’s instructions (Cayman Chemical). Results were calculated according to the standard curve generated by using standards from the kits. Final metabolite concentrations for each sample were normalized to total protein concentration measured by Bradford assay, and values are shown as percent of young 4 mo old mice of the respective genotype, as D2-*Gpnmb*^+^ mice had lower values at all ages.

### Metabolomics

#### Reagents and chemicals

LC-MS grade water was purchased from Honeywell (Riedel-de Hae n; LOT: I174P) and formic acid was purchased from Sigma-Aldrich (LOT: 179247). LiChromasolv hypergrade Acetonitrile (LOT: 10947229817) and 2-propanol (LOT: K50189881818) for LC-MS analysis were purchased from Sigma-Aldrich. The internal lock masses (purine and HP-0921) and tune mix for calibrating the Q-TOF-MS (ESI-low concentration tuning mix) were purchased from Agilent Technologies (USA).

#### Sample preparation

Samples (in 1.5 mL Eppendorf tubes) were kept on dry-ice whilst 150 µL of ice-cold methanol (Optima^T^ LC-MS, Fisher Scientific; LOT: 1862991) was added to each retina sample with a multichannel pipette. Samples were vortexed for 15 seconds prior to sonication in an ice-bath for 15 minutes. Samples were then centrifuged at 12,000 g for 12 minutes at 4 °C. Upon completion, 140 µL of each sample supernatant was added to a separate Ultrafree-MC VV Centrifugal Filter (Sigma-Aldrich; LOT: R7JA29961) and centrifuged for 4 minutes at 8,000 g at original temperature. The supernatant (40 µL each) was transferred to two separate LC vials for ZIC-HILIC and Reversed Phase analysis. Another 40 µL was collected from each sample, and pooled together to represent the quality control (QC). The samples were randomized and after every batch of 12 samples a QC sample was incorporated in order to observe any instrumental drifts, changes in instrument sensitivity, as well as evaluate the %CV of each detected metabolite.

Liquid chromatography-high resolution mass spectrometry (LC-HRMS) experiments were performed on a 1290 Infinity II ultra-high performance liquid chromatography (UHPLC) system coupled to a 6550 iFunnel quadrupole-time of flight (Q-TOF) mass spectrometer equipped with a dual AJS electrospray ionization source (Agilent Technologies). Polar metabolites were separated on a SeQuant ZIC-HILIC (Merck, Darmstadt, Germany) column 100 Å (100 mm × 2.1 mm, 3.5 µm particle size) coupled to a guard column (20 mm × 2.1 mm, 3.5 µm particle size) and an inline-filter. Non-polar metabolites were separated on an EclipsePlus C18 2.1 × 100 mm, 1.8 μm column and guard column (2.1 ×5 mm) and an in-line filter. For the zic-Hilic analysis, mobile phases consisted of 0.1 % formic acid in water with (solvent A) and 0.1 % formic acid in acetonitrile with (solvent B). The elution gradient used was as follows: isocratic step at 95 % B for 1.5 min, 95 % B to 40 % B in 12 mins and maintained at 40 % B for 2 mins, then decreasing to 25 % B at 14.2 min and maintained for 2.8 min, then returned to initial conditions over 1 min, and the column was equilibrated at initial conditions for 7 min. The flow rate was 0.3 mL min-1, injection volume was 3 µL and the column oven was maintained at 25 °C. For the RP analysis, mobile phases consisted of 0.1% formic acid in water (solvent A) and 2-propanol:acetonitrile (90:10, v/v) with 0.1% formic acid (solvent B). The gradient elution was set as follows: isocratic step of 5% B for 3 min, 5 to 30% B in 2 min, then B was increased to 98% in 13.5 min, maintained at 98% B for 1.5 min, returned to initial conditions over 0.5 min, and then held for a further 4.5 min. The flow rate was 0.4 mL/min, injection volume was 3 μL, and the column oven was maintained at 50 °C. Two independent injections were run for positive and negative acquisition modes. The Q-TOF MS system was calibrated and tuned according to the protocols recommended by the manufacturer. Nitrogen (purity > 99.9990%) was used as a sheath gas and drying gas at a flow of 8 L min-1 and 15 L min-1, respectively. The drying and sheath gas temperature were set at 250 °C, with the nebulizer pressure at 35 psi and voltage 3000 V (+/- for positive and negative ionization mode, respectively). The fragmentor voltage was set at 380 V. The acquisition was obtained with a mass range of 50-1200 m/z, where full scan high-resolution data were acquired at three alternating collision energies (0 eV, 10 eV and 30 eV). The data acquisition rate was 6 scans sec-1. Between 16 and 25 min, LC flow was diverted to the waste for the zicHILIC analysis. For further details regarding the acquisition methodology see Naz *et al*. (*34, 35*). Positive and negative raw LC-HRMS files were independently processed with an in-house developed PCDL library for polar and non-polar metabolites using Profinder version B.06 (Agilent Technologies). Metabolites detected in both ionization modes were correlated, and the ones with a larger relative intensity as well as lower %CV were selected. Proper identification of reported compounds (ZIC-HILIC = 52, reversed phase = 33) was assessed by accurate mass and retention time (AMRT) plus fragment identification at two collision energies (10 and 30 eV) using an in-house generated database. A separate PCDL library was created to screen for isoleucine putatively. Metabolomics data was analyzed using MetaboAnalyst and R using the 86 identified metabolites as a background list for enrichment (*36, 37*). Metabolite identities were confirmed according to Level 1 of the Metabolomics Standards Initiative (*11*).

### mTOR phospho antibody array

mTOR phospho antibody arrays were performed on *n* = 3 per condition whole homogenized retina from 9 month of age D2 and D2-*Gpnmb*^+^ mice following the manufacturer’s instructions (Full Moon BioSystems; PMT138). Data was analyzed in R using limma (linear models for microarray data, designed for microarray analysis but suitable for other data types) (*38-40*).

### RNA-sequencing

RNA-sequencing data was accessed through the Gene Expression Omnibus (accession number GSE90654) and analyzed as previously described in R (*10*).

### Statistical analysis

The sample size (number of eyes, *n*) is shown in each figure legend. Graphing and statistical analysis was performed in R. *Student’s t* test was used for pairwise analysis in quantitative plots, and *Fisher’s exact* test for optic nerve analysis. Where multiple groups were compared, a *one-way ANOVA* was employed, with *post-hoc Tukey-Kramer* test. For boxplots, center hinge represents the median, and the upper and lower hinges represent the first and third quartiles, whiskers represent 1.5 X interquartile range, values beyond the whiskers are plotted as outliers. Unless otherwise stated * = *P* < 0.05, ** = *P* < 0.01, *** = *P* < 0.001.

## Supplementary figure legends

**Fig. S1. Pyruvate dysregulation during early glaucoma pathogenesis.** (**A**) Pathway analysis of RNA-sequencing data from pre-glaucomatous D2 RGCs demonstrates significant enrichment of glucose and pyruvate related metabolic pathways. D2 Groups 1-4 represent increasing disease-status at the transcriptomic level as identified in Williams *et al.* 2017 (*10*). (**B and C**) Retinal pyruvate declines following chronic IOP-elevation in D2 eyes without age-related changes (as demonstrated in control age-matched retinas) (*n* > 24/group). (**D and E**) MPC1 and (**F and G**) MPC2 are robustly expressed in the ganglion cell layer (RGC somas) and inner plexiform layer (RGC dendrites) in mouse, rat, marmoset, and human retina as assessed by immunofluorescence and immunohistochemistry. GCL = ganglion cell layer, IPL = inner plexiform layer, INL = inner nuclear layer. Black arrows = RGCs with high expression of MPC1 and MPC2.

**Fig. S2. Pyruvate treatment protects from glaucoma-like stresses in retinal neurons.** Pyruvate protects from RGC degeneration following glucose deprivation (**A and B**; *n* = 24/group) and from *t*-butyl hydroperoxide (t-bH) induced oxidative stress (as assessed by MTT viability assay, **C-E**; *n* = 8/group). Addition of 10 µM 4CIN (a monocarboxylate transporter inhibitor) demonstrates that pyruvate protects as a metabolic substrate and is imported into RGCs. For **E**, t-bH = 100 μM.

**Fig. S3. Pyruvate treatment protects from RGC axotomy.** Pyruvate protects from cell loss (**A and B**) and confers partial protection to pre-apoptotic nucleus shrinkage (**C**) following axotomy in the mouse retina explant axotomy model (*n* = 8/group).

**Fig. S4. Iris disease and IOP elevation in treated mice.** Treatment with NAM and pyruvate did not prevent iris degeneration, pigment dispersion (**A**; 12 mo), nor IOP elevation in D2 mice (*P* > 0.05 across all comparisons to untreated) (**B**). The age of onset, rate of progression, and severity of iris damage was similar between groups. After eight weeks of treatment, rapamycin caused bleeding and haziness in the anterior of the eye and grossly increased the amount of iris depigmentation compared to untreated mice (**C**; 8 mo). High IOP was observed in rapamycin treated mice at expected ages for D2 mice (*P* > 0.05 D2 treated to untreated control) (**D**). For dot plots, the diamond (*red*) represents the 95% confidence interval of the mean (centerline).

**Fig. S5. Metabolic and transcriptomic dysregulation of GSH and NAD dependent pathways.** Schematic pathway of glutathione metabolism (**A**) and the NAD salvage pathway (**B**) integrating metabolomic and RNA-sequencing data.

**Fig. S6. Pyruvate treatment rescues many IOP-dependent changes in the retinal metabolome and generates a new metabolic state.** (**A**) Whole retina samples underwent untargeted metabolomic profiling. HC (*Spearman’s*) defined metabolomically distinct groups between retinas from control glaucoma and pyruvate treated retinas (*n* = 10/group). (**B**) Pyruvate treated D2 mice had differences in 27 metabolites compared to control glaucoma retinas, including lower levels of glucose and higher levels of NAM. (**C**) KEGG and (**D**) SMPDB pathway analysis of metabolites demonstrates significant pathway enrichment of pathways related to glucose and pyruvate metabolism and NAD metabolism.

**Fig. S7. Combination therapy with pyruvate and NAM.** (**A and B**) Pyruvate and NAM combination therapy protected from optic nerve degeneration as assessed by PPD staining and was more protective than either treatment alone (*n* > 60/group). Green, no or early damage [<5% axon loss; no or early (NOE)]; yellow, moderate damage (∼30% axon loss; MOD); red, severe (>50% axon loss; SEV) damage. (**C**) Counts and (**D**) optic nerve areas from D2 mice demonstrate a robust axon saving in mice protected by each treatment (*n* = 10/group). (**E**) Pyruvate and NAM combination treatment protected from loss in PERG amplitude (*n* > 24/group).

## Supplementary tables

**Table S1. Metabolomics contrasts.** *See attached documents.*

**Table S2. Metabolomics pathway analysis.** *See attached documents.*

---

**Table S3.**
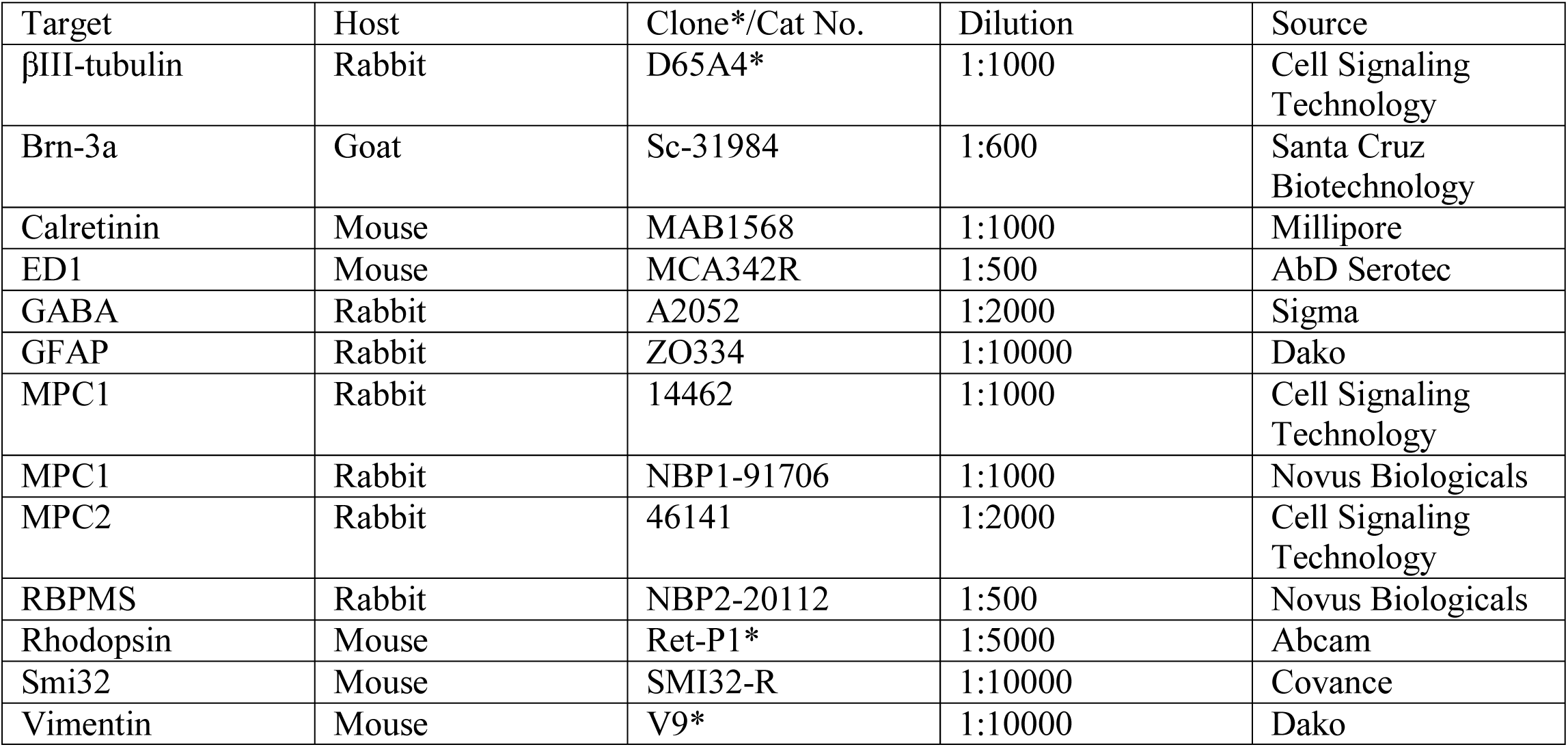
Antibodies used

## References

1. P. A. Williams, J. M. Harder, S. W. M. John, Glaucoma as a Metabolic Optic Neuropathy: Making the Case for Nicotinamide Treatment in Glaucoma. J Glaucoma, (2017).

2. D. M. Inman, M. Harun-Or-Rashid, Metabolic Vulnerability in the Neurodegenerative Disease Glaucoma. Front Neurosci 11, 146 (2017).

3. L. R. Gray, S. C. Tompkins, E. B. Taylor, Regulation of pyruvate metabolism and human disease. Cell Mol Life Sci 71, 2577–2604 (2014).

4. Y. Zilberter, O. Gubkina, A. I. Ivanov, A unique array of neuroprotective effects of pyruvate in neuropathology. Front Neurosci 9, 17 (2015).

5. S. A. Andrabi et al., Poly(ADP-ribose) polymerase-dependent energy depletion occurs through inhibition of glycolysis. Proc Natl Acad Sci U S A 111, 10209–10214 (2014).

6. G. R. Howell et al., Radiation treatment inhibits monocyte entry into the optic nerve head and prevents neuronal damage in a mouse model of glaucoma. J Clin Invest 122, 1246–1261 (2012).

7. P. A. Williams et al., Inhibition of monocyte-like cell extravasation protects from neurodegeneration in DBA/2J glaucoma. Mol Neurodegener 14, 6 (2019).

8. P. A. Williams, N. Marsh-Armstrong, G. R. Howell, L. I. I. o. A. a. G. N. Participants, Neuroinflammation in glaucoma: A new opportunity. Exp Eye Res 157, 20–27 (2017).

9. V. Chrysostomou, F. Rezania, I. A. Trounce, J. G. Crowston, Oxidative stress and mitochondrial dysfunction in glaucoma. Curr Opin Pharmacol 13, 12–15 (2013).

10. P. A. Williams et al., Vitamin B3 modulates mitochondrial vulnerability and prevents glaucoma in aged mice. Science 355, 756–760 (2017).

11. L. W. Sumner et al., Proposed minimum reporting standards for chemical analysis Chemical Analysis Working Group (CAWG) Metabolomics Standards Initiative (MSI). Metabolomics 3, 211–221 (2007).

12. Y. An et al., Evidence for brain glucose dysregulation in Alzheimer’s disease. Alzheimers Dement 14, 318–329 (2018).

13. J. Shi et al., Review: Traumatic brain injury and hyperglycemia, a potentially modifiable risk factor. Oncotarget 7, 71052–71061 (2016).

14. S. Roy, D. Kim, A. Sankaramoorthy, Mitochondrial Structural Changes in the Pathogenesis of Diabetic Retinopathy. J Clin Med 8, (2019).

15. A. Dlasková et al., 3D super-resolution microscopy reflects mitochondrial cristae alternations and mtDNA nucleoid size and distribution. Biochim Biophys Acta Bioenerg 1859, 829–844 (2018).

16. Z. Qi et al., Characterization of susceptibility of inbred mouse strains to diabetic nephropathy. Diabetes 54, 2628–2637 (2005).

17. I. Soto et al., DBA/2J mice are susceptible to diabetic nephropathy and diabetic exacerbation of IOP elevation. PLoS One 9, e107291 (2014).

18. J. Kouassi Nzoughet et al., Nicotinamide Deficiency in Primary Open-Angle Glaucoma. Invest Ophthalmol Vis Sci 60, 2509–2514 (2019).

19. P. A. Williams, J. M. Harder, B. H. Cardozo, N. E. Foxworth, S. W. M. John, Nicotinamide treatment robustly protects from inherited mouse glaucoma. Commun Integr Biol 11, e1356956 (2018).

20. P. Williams et al., Nicotinamide and WLDS act together to prevent neurodegeneration in glaucoma. Frontiers in Neuroscience 11, (2017).

21. A. H. Jassim et al., Higher Reliance on Glycolysis Limits Glycolytic Responsiveness in Degenerating Glaucomatous Optic Nerve. Mol Neurobiol 56, 7097–7112 (2019).

22. M. Knip et al., Safety of high-dose nicotinamide: a review. Diabetologia 43, 1337–1345 (2000).

23. R. A. Saxton, D. M. Sabatini, mTOR Signaling in Growth, Metabolism, and Disease. Cell 168, 960–976 (2017).

24. X. Zheng et al., Alleviation of neuronal energy deficiency by mTOR inhibition as a treatment for mitochondria-related neurodegeneration. Elife 5, (2016).

25. S. W. John et al., Essential iris atrophy, pigment dispersion, and glaucoma in DBA/2J mice. Invest Ophthalmol Vis Sci 39, 951–962 (1998).

26. H. Levkovitch-Verbin et al., Translimbal laser photocoagulation to the trabecular meshwork as a model of glaucoma in rats. Invest Ophthalmol Vis Sci 43, 402–410 (2002).

27. M. Saleh, M. Nagaraju, V. Porciatti, Longitudinal evaluation of retinal ganglion cell function and IOP in the DBA/2J mouse model of glaucoma. Invest Ophthalmol Vis Sci 48, 4564–4572 (2007).

28. T. H. Chou, J. Bohorquez, J. Toft-Nielsen, O. Ozdamar, V. Porciatti, Robust mouse pattern electroretinograms derived simultaneously from each eye using a common snout electrode. Invest Ophthalmol Vis Sci 55, 2469–2475 (2014).

29. R. Smith, S. John, P. Nishina, J. Sundberg, *Systematic evaluation of the mouse eye. Anatomy, pathology and biomethods*. (CRC Press, Boca Raton, 2002).

30. K. Zarei et al., Automated Axon Counting in Rodent Optic Nerve Sections with AxonJ. Sci Rep 6, 26559 (2016).

31. G. Chidlow, J. P. Wood, B. Knoops, R. J. Casson, Expression and distribution of peroxiredoxins in the retina and optic nerve. Brain Struct Funct 221, 3903–3925 (2016).

32. J. P. Wood, T. Mammone, G. Chidlow, T. Greenwell, R. J. Casson, Mitochondrial inhibition in rat retinal cell cultures as a model of metabolic compromise: mechanisms of injury and neuroprotection. Invest Ophthalmol Vis Sci 53, 4897–4909 (2012).

33. L. Peng et al., Glutamine as an energy substrate in cultured neurons during glucose deprivation. J Neurosci Res 85, 3480–3486 (2007).

34. S. Naz et al., Development of a Liquid Chromatography-High Resolution Mass Spectrometry Metabolomics Method with High Specificity for Metabolite Identification Using All Ion Fragmentation Acquisition. Anal Chem 89, 7933–7942 (2017).

35. E. Daskalaki, N. J. Pillon, A. Krook, C. E. Wheelock, A. Checa, The influence of culture media upon observed cell secretome metabolite profiles: The balance between cell viability and data interpretability. Anal Chim Acta 1037, 338–350 (2018).

36. J. Chong et al., MetaboAnalyst 4.0: towards more transparent and integrative metabolomics analysis. Nucleic Acids Res 46, W486–W494 (2018).

37. J. Chong, D. S. Wishart, J. Xia, Using MetaboAnalyst 4.0 for Comprehensive and Integrative Metabolomics Data Analysis. Curr Protoc Bioinformatics 68, e86 (2019).

38. M. E. Ritchie et al., Empirical array quality weights in the analysis of microarray data. BMC Bioinformatics 7, 261 (2006).

39. M. E. Ritchie et al., limma powers differential expression analyses for RNA-sequencing and microarray studies. Nucleic Acids Res 43, e47 (2015).

40. G. K. Smyth, J. Michaud, H. S. Scott, Use of within-array replicate spots for assessing differential expression in microarray experiments. Bioinformatics 21, 2067–2075 (2005)

